# A bright and high-performance genetically encoded Ca^2+^ indicator based on mNeonGreen fluorescent protein

**DOI:** 10.1101/2020.01.16.909291

**Authors:** Landon Zarowny, Abhi Aggarwal, Virginia M.S. Rutten, Ilya Kolb, The GENIE Project, Ronak Patel, Hsin-Yi Huang, Yu-Fen Chang, Tiffany Phan, Richard Kanyo, Misha Ahrens, W. Ted Allison, Kaspar Podgorski, Robert E. Campbell

**Affiliations:** Department of Chemistry, University of Alberta, Edmonton, Alberta, Canada; Janelia Research Campus, Howard Hughes Medical Institute, Ashburn, Virginia, United States; Gatsby Computational Neuroscience Unit, UCL, London, UK; LumiSTAR Biotechnology, Inc. National Biotechnology Research Park, Taipei City 115, Taiwan; Department of Biological Sciences, University of Alberta, Edmonton, Alberta, Canada; Department of Chemistry, Graduate School of Science, The University of Tokyo, Tokyo, Japan

## Abstract

Genetically encodable calcium ion (Ca^2+^) indicators (GECIs) based on green fluorescent proteins (GFP) are powerful tools for imaging of cell signaling and neural activity in model organisms. Following almost two decades of steady improvements in the *Aequorea victoria* GFP (avGFP)-based GCaMP series of GECIs, the performance of the most recent generation (i.e., GCaMP7) may have reached its practical limit due to the inherent properties of GFP. In an effort to sustain the steady progression towards ever-improved GECIs, we undertook the development of a new GECI based on the bright monomeric GFP, mNeonGreen (mNG). The resulting indicator, mNG-GECO1, is 60% brighter than GCaMP6s *in vitro* and provides comparable performance as demonstrated by imaging Ca^2+^ dynamics in cultured cells, primary neurons, and *in vivo* in larval zebrafish. These results suggest that mNG-GECO1 is a promising next-generation GECI that could inherit the mantle of GCaMP and allow the steady improvement of GECIs to continue for generations to come.

## Introduction

Genetically encodable calcium ion (Ca^2+^) indicators (GECIs) are a class of single fluorescent protein (FP)-based biosensors that are powerful tools for the visualization of Ca^2+^ concentration dynamics both *in vitro* and *in vivo*^1, 2, 3^. As they are genetically encoded, GECI expression can be genetically targeted to specific cell types or subcellularly localized to specific organelles. Furthermore, their negligible cellular toxicity, minimal perturbation of endogenous cellular functions, and biological turnover, make them ideal for long-term imaging experiments^4^. The Ca^2+^-dependent fluorescent response of GECIs is routinely used as a proxy for neuronal activity due to the transient changes in Ca^2+^ concentration that accompany action potentials^5, 6, 7, 8^. GECIs have facilitated the optical recording of thousands of neurons simultaneously in the surgically exposed brains of mice^9^. Despite their widespread use by the scientific community, there are some properties of GECIs that could be further improved. These properties include faster Ca^2+^ response kinetics, higher fluorescent molecular brightness, and minimized contribution to Ca^2+^ buffering. Some GECIs have shown aggregation in neurons, and some of the most highly optimized GECIs have been demonstrated to cause aberrant cortical activity in murine models^10, 11, 12^.

An important issue that is common to all GECIs is their intrinsic Ca^2+^ buffering capacity. The Ca^2+^ binding domains of GECIs (calmodulin (CaM) or troponin C (TnC)) act as Ca^2+^ buffers within the cell and must necessarily compete with endogenous proteins for binding to Ca^2+^ (Refs. 13, 14, 15, 16). Comprehensive investigations of this phenomenon are limited, but a few reports have indicated abnormal morphology and behavior of neurons after long term or high expression of GCaMPs^17^. Ca^2+^ buffering and competition for CaM binding sites have been proposed as possible causes. One solution to the Ca^2+^ buffering phenomenon is to reduce the reporter protein expression, leading to a lower concentration of GECI and reduced buffering capacity. However, reduced expression requires increased intensity of excitation light to achieve an equivalent fluorescent signal, which can lead to increased phototoxicity and photobleaching. Another solution is to reduce the number of Ca^2+^ binding sites like that in the TnC-based GECIs, NTnC^18^ and YTnC^19^. Unfortunately, these indicators have relatively low fluorescence response (ΔF/F_min_ ~ 1 for NTnC and ~ 10.6 for YTnC) compared to the recent GCaMP7 variants (ΔF/F_min_ ~ 21 to 145)^20^. Another possible solution is to develop GECIs with increased brightness such that they could be expressed at a lower concentration while retaining a similar fluorescent intensity with similar intensity of excitation light.

Further increasing the brightness of GECIs, while retaining high performance comparable to the most recent generation of indicators, would provide improved tools for optical imaging of neuronal activity and decrease the occurrence of experimental artifacts resulting from Ca^2+^ buffering and indicator overexpression^17^. Our efforts to realize this advance were inspired, in part, by the advent of a bright and monomeric engineered version of GFP from *Branchiostoma lanceolatum*, mNeonGreen (mNG)^21^. Due to its high brightness and its excellent performance as a subcellular localization tag^21^, mNG is an exceptionally promising starting point from which to develop a brighter GECI.

Here we introduce an <u>mNG</u>-based genetically <u>e</u>ncodable <u>C</u>a^2+^ indicator for <u>o</u>ptical imaging (mNG-GECO1) that exceeds the brightness of all variants in the GCaMP series while providing performance that is comparable to the latest generation GCaMP variants. Key design differences between mNG-GECO1 and the GCaMP series include the GFP portion (mNG versus avGFP) and the protein topology (non-circularly permuted mNG versus circularly permuted avGFP).

## Results and Discussion

### Rational engineering and iterative directed evolution of mNG-GECO1

We used a combination of rational design, linker sequence optimization, and directed evolution to develop mNG-GECO1 (**Supplementary Fig. 1**). Starting from an unpublished topological variant of REX-GECO1^22^, we used PCR to produce a fragment containing CaM linked to the RS20 peptide with a short linker (**Fig. 1a**). Insertion of this PCR fragment into the mNG gene between residues 136 and 137 (numbering as in PDB ID 5LTR)^23^ resulted in a green fluorescent indicator prototype which we named mNG-GECO0.1 (**Fig. 1**). For the remainder of this manuscript, amino acids will be numbered as in the sequence alignment provided as **Supplementary Fig. 2**. mNG-GECO0.1 had a minimal response to Ca^2+^ (ΔF/F_min_ = 0.3), but we anticipated that optimization of the sequence around the insertion site would yield a suitable template for directed evolution. Indeed, we found that deletion of Ala146, the residue immediately preceding the insertion of the Ca^2+^ sensing domain, substantially improved the response to Ca^2+^ (mNG-GECO0.2; ΔF/F_min_ ~ 2).

**Figure 1.**
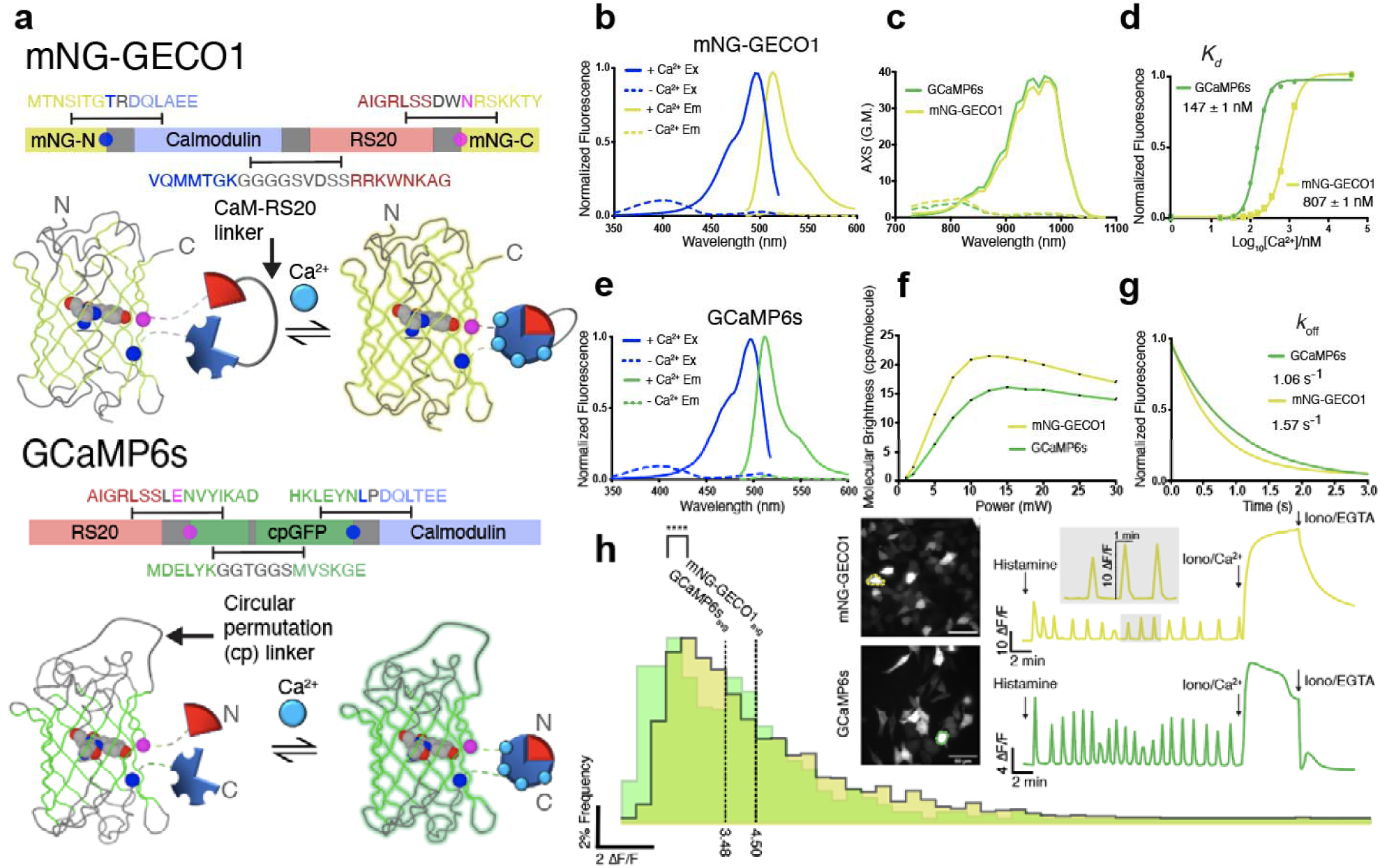
Topology and *in vitro* characterization of mNG-GECO1 and GCaMP6s. **a** Topology of non-circularly permuted mNG-GECO1 and circularly permuted GCaMP6s. Linker regions are shown in grey and the two residues that flank the insertion site (residue 136 of mNG in blue and residue 139 in magenta; numbering as in PDB ID 5LTR)^23^ are shown as circles on both the protein structure and gene schematics. The Ca^2+^ responsive domains are shaded light blue for CaM and light red for RS20. **b**,**e** Excitation and emission spectra for each indicator. **c** 2-photon cross section for each indicator in Ca^2+^ saturated or Ca^2+^ free states. **d** Ca^2+^ titration for GCaMP6s (*K*_d_ = 147 ± 1 nM) and mNG-GECO1 (807 ± 1 nM). **f** Dependence of two-photon molecular brightness on excitation power intervals. **g** Stop-flow kinetics for each indicator showing mNG-GECO1 (*k*_off_ 1.57 s^−1^) and GCaMP6s (*k*_off_ 1.06 s^−1^). **h** Characterization of histamine induced Ca^2+^ oscillations in HeLa cells with representative traces inset.

Starting from mNG-GECO0.2, we began a process of iterative directed evolution which involved screening of libraries created from error-prone PCR or site saturation mutagenesis to identify variants with increased brightness and increased response to Ca^2+^. In our primary library screen, we used a fluorescent colony screening system equipped with excitation and emission filters appropriate for imaging of green fluorescence^24^. Bright colonies were picked and cultured overnight in liquid media. A secondary screen for Ca^2+^ sensitivity was performed the next day using detergent-extracted bacterial lysate. The fluorescence of the lysate for each variant was measured in Ca^2+^ chelating buffer (30 mM MOPS, 100 mM KCl, 10 mM EGTA, pH 7.3), and subsequently in Ca^2+^ saturating buffer (30 mM MOPS, 100 mM KCl, 10 mM Ca^2+^, pH 7.3). Dividing the Ca^2+^ saturated fluorescence by the Ca^2+^ free fluorescence provided an approximate but robust measure of each indicator variant’s response to Ca^2+^. For each round of screening the plasmids were isolated for the 6-10 most promising variants and sent for sequencing. The pool of these most promising variants was used as the template for the next round of library creation and directed evolution.

Following 7 rounds of iterative directed evolution, *E. coli* colonies harboring mNG-GECO0.7 were brightly fluorescent after overnight incubation. However, the Ca^2+^ response of mNG-GECO0.7 was remained relatively low (ΔF/F_min_ ~ 5), relative to recent generation GCaMP variants. We anticipated that optimization of the linkers connecting mNG to the CaM-RS20 domain (mNG-CaM linker and RS20-mNG linker) could lead to the identification of variants with improved responses. To optimize these linker regions, we used site saturation mutagenesis to produce libraries of all 20 amino acids within the 3 residues connecting mNG to CaM. Individual libraries of Leu133, Thr134, and Ala135 were randomized to all 20 amino acids. If a beneficial mutation was found, the process was repeated for the remaining amino acids until these libraries were exhausted. By screening of these libraries, we identified two mutations of the linker region between mNG barrel and CaM: Ala145Gly and Leu143Ile. This variant, mNG-GECO0.9, had a ΔF/F_min_ ~ 12 as measured *in vitro*.

Following optimization of the mNG-CaM linker, multiple site saturation libraries were created, using the same methodology as the mNG-CaM linker, for the RS20-mNG linker region (residues Glu323, Trp324, Cys325 and Arg326). Screening of these libraries led to the identification of a particularly bright variant with a Cys325Asn mutation. This variant, designated mNG-GECO0.9.1, is brighter than mNG-GECO0.9 but has a decreased response to Ca^2+^ of ΔF/F_min_ = 3.5. In an effort to improve the performance of mNG-GECO0.9.1, we applied site saturations to positions previously found to be mutated during directed evolution. Screening of these libraries for variants with increased brightness and higher ΔF/F_min_ led to the identification of a variant with Asp206Gly, Phe209Leu, Pro263Phe, Lys265Ser, Thr346Ile and the reversion of Gly152Glu. This variant was designated as mNG-GECO1. A notable observation from the directed evolution efforts is the minimal number of mutations in the mNG domain. Only 2 mutations (Lys128Glu and Thr346Ile) were outside the β-strand in which the Ca^2+^ sensing domain was inserted. In contrast, 3 mutations were localized to the β-strand surrounding the sensing domain insertion site (Leu143Ile/Ala145Gly/Cys325Asn) and 7 mutations were localized to the CaM domain (Thr151Ala/Thr180Cys/Asp206Gly/Phe209Leu/Pro263Phe/Lys265Ser/Ala293Gly).

### *In vitro* characterization of mNG-GECO1

We characterized mNG-GECO1, in parallel with GCaMP6s, for direct comparison of biophysical properties measured under identical conditions (**Supplementary Table 1).** We found that the excitation (ex) and emission (em) maxima of the Ca^2+^ saturated states to be 497 nm (ex) and 512 nm (em) for GCaMP6s and 496 nm (ex) and 513 nm (em) for mNG-GECO1 (**Fig. 1**). The *in vitro* Ca^2+^ response of mNG-GECO1 (ΔF/F_min_ = 35) was similar to that of GCaMP6s when tested in parallel (ΔF/F_min_ = 39). The *K*_d_ of mNG-GECO1 (810 nM) is substantially higher than that of GCaMP6s (147 nM). This increase in *K*_d_ is consistent with the faster *k*_off_ kinetics of mNG-GECO1 (*k*_off_ = 1.57 +/− 0.01 s^−1^) relative to GCaMP6s (*k*_off_ = 1.06 +/− 0.01 s^−1^). There was no noticeable difference observed between the *k*_on_ kinetics of mNG-GECO1 and GCaMP6s with varying concentrations of Ca^2+^ (**Supplementary Fig. 3**).

In the Ca^2+^ bound state, mNG-GECO1 has an extinction coefficient of 102,000 M^−1^cm^−1^ and quantum yield of 0.69, giving it an overall brightness (= EC * QY) of 70. This value is similar to the value of 77 previously reported for NTnC^18^ and 78% of the brightness of mNG itself (measured by us to be 112,000 M^−1^cm^−1^ * 0.8 = 90) (**Supplementary Fig. 4**). Under two-photon excitation conditions, both mNG-GECO1 and GCaMP6s have a maximal two-photon cross section at ~ 970 nm and similar action cross-section (AXS) values of 37.22 GM for mNG-GECO1 and 38.81 GM for GCaMP6s. However, due to its higher brightness at the single molecule level, the molecular brightness of mNG-GECO1 (21.3) is higher than that of GCaMP6s (16.1) at 15 mW power. Overall, these data indicate the mNG-GECO1 has excellent one-photon and two-photon excitation properties *in vitro*.

### In vitro characterization in cultured cells and dissociated neurons

To compare the performance of mNG-GECO1 and GCaMP6s in cultured cells, we transfected HeLa cells with mNG-GECO1 in a pcDNA vector (CMV promoter) in parallel with pGP-CMV-GCaMP6s. Using a previously reported protocol^25^, Ca^2+^ oscillations were induced by treatment with histamine and fluorescence images were acquired every 10 seconds for 20 minutes. From the intensity versus time data for each cell, ΔF/F_0_ for all oscillations of ΔF/F_0_ > 0.5 were extracted using a Matlab script. Using these extracted ΔF/F_0_ values, average ΔF/F_0_ for all oscillations and maximum ΔF/F_0_, was computed. The average maximum ΔF/F_0_ was calculated by averaging the maximum ΔF/F_0_ from each responding cell. In parallel experiments, mNG-GECO1 had an average ΔF/F_0_ = 4.50 ± 2.96 compared to GCaMP6s’ ΔF/F_0_ = 3.48 ± 2.40 (**Fig. 1h**). The maximum ΔF/F_0_ was 16.8 ± 10.5 for mNG-GECO1 and 12.8 ± 6.11 for GCaMP6s. At the end of the 20-minute imaging experiment, the cells were treated with ionomycin/Ca^2+^ to saturate the indicators and induce a fluorescent maximum and then with Ca^2+^ chelator EGTA/ionomycin to deplete Ca^2+^ and produce a fluorescent minimum. For these treatments, ΔF/F_min_ = 48.8 ± 15.1 for mNG-GECO1 and ΔF/F_min_ = 16.7 ± 5.2 for GCaMP6s. These results were obtained from a data set of 137 responding cells with 1624 individual oscillations for mNG-GECO1 and 99 responding cells with 687 individual oscillations for GCaMP6s (**Supplementary Table 2**).

We next characterized the performance of mNG-GECO1 in dissociated rat cortical neurons alongside GCaMP series indicators GCaMP6s, jGCaMP7s, jGCaMP7b, jGCaMP7c, and jGCaMP7f (**Fig. 2**). Field stimulated neurons expressing mNG-GECO1 had a single action potential (AP) ΔF/F_0_ = 0.19 ± 0.04, slightly lower than that of GCaMP6s (ΔF/F_0_ = 0.27 ± 0.09, **Fig. 2a**). For 10 APs, performance of mNG-GECO1 was approximately twofold lower than GCaMP6s, with ΔF/F_0_ of 1.5 ± 0.19 and 3.1 ± 0.26 for mNG-GECO1 and GCaMP6s, respectively (**Fig. 2b**). At 160 APs, mNG-GECO1 has a ΔF/F_0_ of 6.5 ± 0.8, slightly lower than GCaMP6s’s ΔF/F_0_ of 9.0 ± 0.47 (**Fig. 2c**). The baseline brightness of mNG-GECO1 (1,374 ± 31 AU) was comparable to the baseline brightness of GCaMP6s (1,302 ± 6 AU) and jGCaMP7s (1,397 ± 11 AU) (**Fig. 2e**). The signal-to-noise ratio (SNR) of mNG-GECO1 and GCaMP6s are comparable for 1 and 3 AP’s (**Fig. 2f**). For 3 AP stimulation, mNG-GECO1 exhibited a half rise time of 49 ± 1 ms and half decay time of 582 ± 12 ms. Under the same conditions, GCaMP6s exhibited a half rise time of 65 ± 2 ms and a half decay time of 1,000 ± 36 ms. Field stimulated neuron data is summarized in **Supplementary Table 3**. The overall data in cultured neuron suggest that the mNG-GECO1 sensor is comparable in signal, kinetics, and baseline brightness to the GCaMP6s sensor.

**Figure 2.**
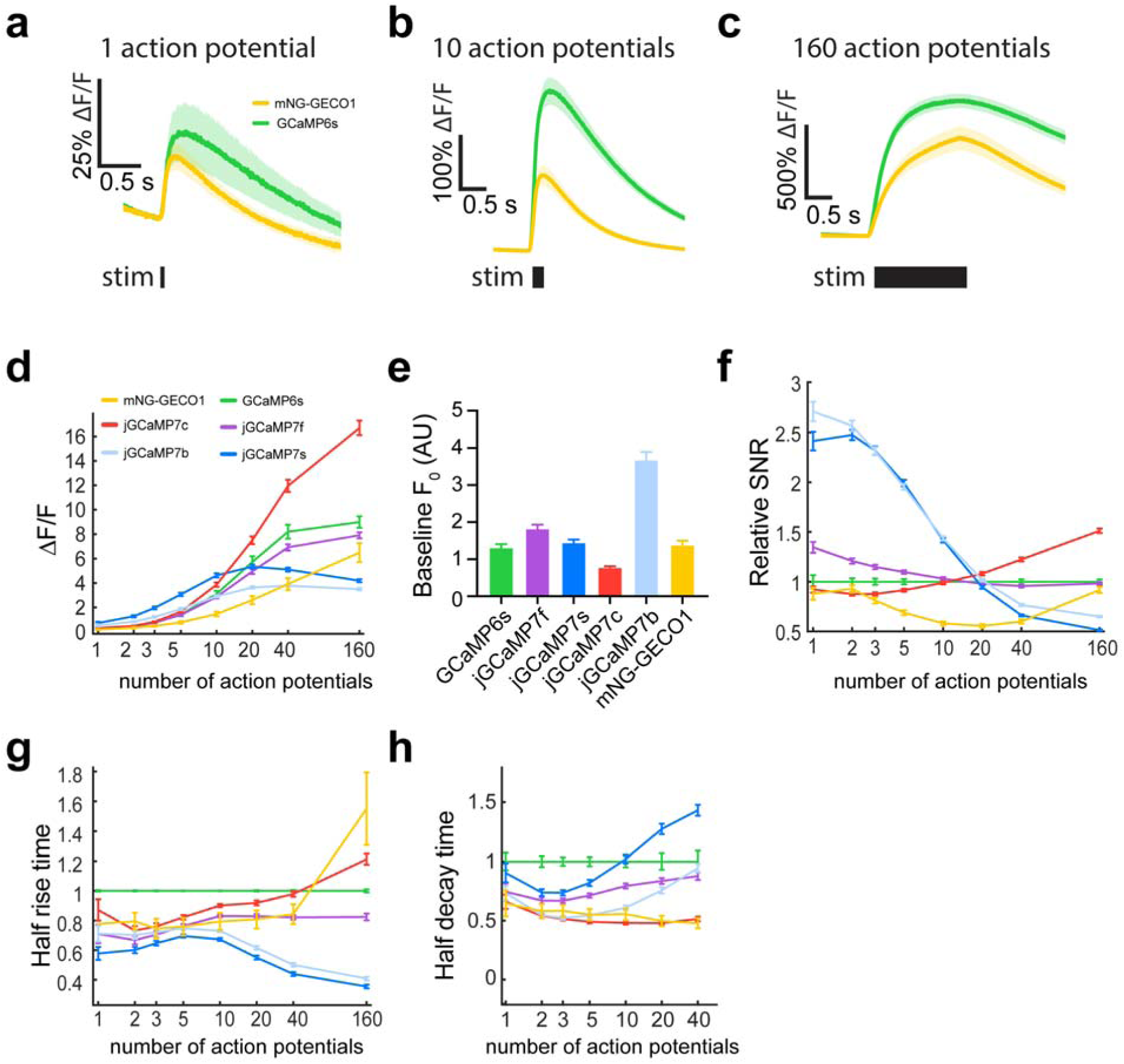
Characterization of mNG-GECO1 and GCaMP series indicators in dissociated rat hippocampal neurons. **a-c** Average responses to 1, 10, and 160 action potentials for mNG-GECO1 and GCaMP6s. Shaded areas correspond to s.e.m. for each trace. **d** Response amplitude ΔF/F_0_ for mNG-GECO1 and the GCaMP series of indicators in response to 1, 2, 3, 5, 10, 20, 40, and 160 action potentials. Data are presented normalized to ΔF/F_0_ of GCaMP6s. **e** Baseline brightness for each indicator, defined as the mean raw fluorescence intensity of all neurons prior to the stimulus. **f** Relative SNR, defined as the peak raw fluorescence divided by the signal standard deviation prior to the stimulus, normalized to SNR of GCaMP6s. **g** Half-rise time normalized to GCaMP6s. **h** Half-decay time normalized to GCaMP6s. The 160 action potential measurement was omitted because fluorescence levels generally did not return to baseline over the imaging period. For **a**-**h**, mNG-GECO1: 621 neurons, 15 wells; GCaMP6s: 937 neurons, 17 wells; jGCaMP7c: 2,551 neurons, 44 wells; jGCaMP7b: 2,339 neurons, 47 wells; jGCaMP7f: 2,585 neurons, 48 wells; and jGCaMP7s: 2,249 neurons, 47 wells. Data in **d**-**h** shown as mean ± s.e.m., see **Supplementary Table 3** for analyzed data.

### *In vivo* evaluation of mNG-GECO1

To evaluate mNG-GECO1 for *in vivo* expression in zebrafish neurons, we used a Tol2 transposase transgenesis system to deliver mNG-GECO1 or GCaMP6s under a pan-neuronal Elavl3 promoter into zebrafish embryos^26^. We tracked expression of mNG-GECO1 over several days to evaluate the viability of transgenic fish (**Supplementary Fig. 5**). We found no obvious morphological anomalies during larval development stage of zebrafish expressing mNG-GECO1 or GCaMP6s.

To evaluate the relative performance of mNG-GECO1 and GCaMP6s for imaging of neuronal activity in zebrafish larvae, we used the same transgenesis protocol to produce *Casper* zebrafish lines expressing each indicator (**Fig. 3**). Prior to imaging, 5-6 days post fertilization *Casper* fish expressing the sensors were immobilized with bungarotoxin (1 mg/mL) for 30 seconds followed by a 10 minute incubation in the convulsant 80 mM 4-aminopyridine (4-AP). The fish were then placed in low melting agar and immersed in a solution of 4-AP (80 mM). Imaging consisted of 5 minute intervals of the hindbrain or midbrain at a recording rate of 3 Hz. For each indicator, 5 fish were imaged under 6 different field of views resulting in 834 and 1280 individual cells for mNG-GECO1 and GCaMP6s, respectively (**Supplementary Fig. 6**). The resulting data was evaluated using the Suite2p package (https://github.com/MouseLand/suite2p)^27^. We found that mNG-GECO1 had a maximum ΔF/F_0_ for each cell of 3.09 ± 0.08 compared to 4.56 ± 0.11 for GCaMP6s (**Fig. 3c**). The baseline fluorescence of GCaMP6s was higher compared to mNG-GECO1 (0.95 ± 0.03 vs 1.41 ± 0.05 AU, respectively) (**Fig. 3d**). However, the signal-to-noise ratio (SNR), which was computed by dividing the ΔF/F_0_ by the raw standard deviation of each cell in six field of views, was higher for mNG-GECO1 (SNR = 6.63 ± 0.07) than GCaMP6s (SNR = 5.25 ± 0.04) (**Fig. 3e**). We also found that mNG-GECO1 had a slower decay time (faster *k*_off_ kinetics) compared to GCaMP6s (1.98 ± 0.12 s^−1^ vs 3.00 ± 0.12 s^−1^, respectively) (**Fig. 3f**). The overall data in zebrafish neurons suggest that mNG-GECO1 is comparable in signal-to-noise ratio, kinetics, and baseline brightness to the GCaMP6s sensor (**Supplementary Table 4**).

**Figure 3.**
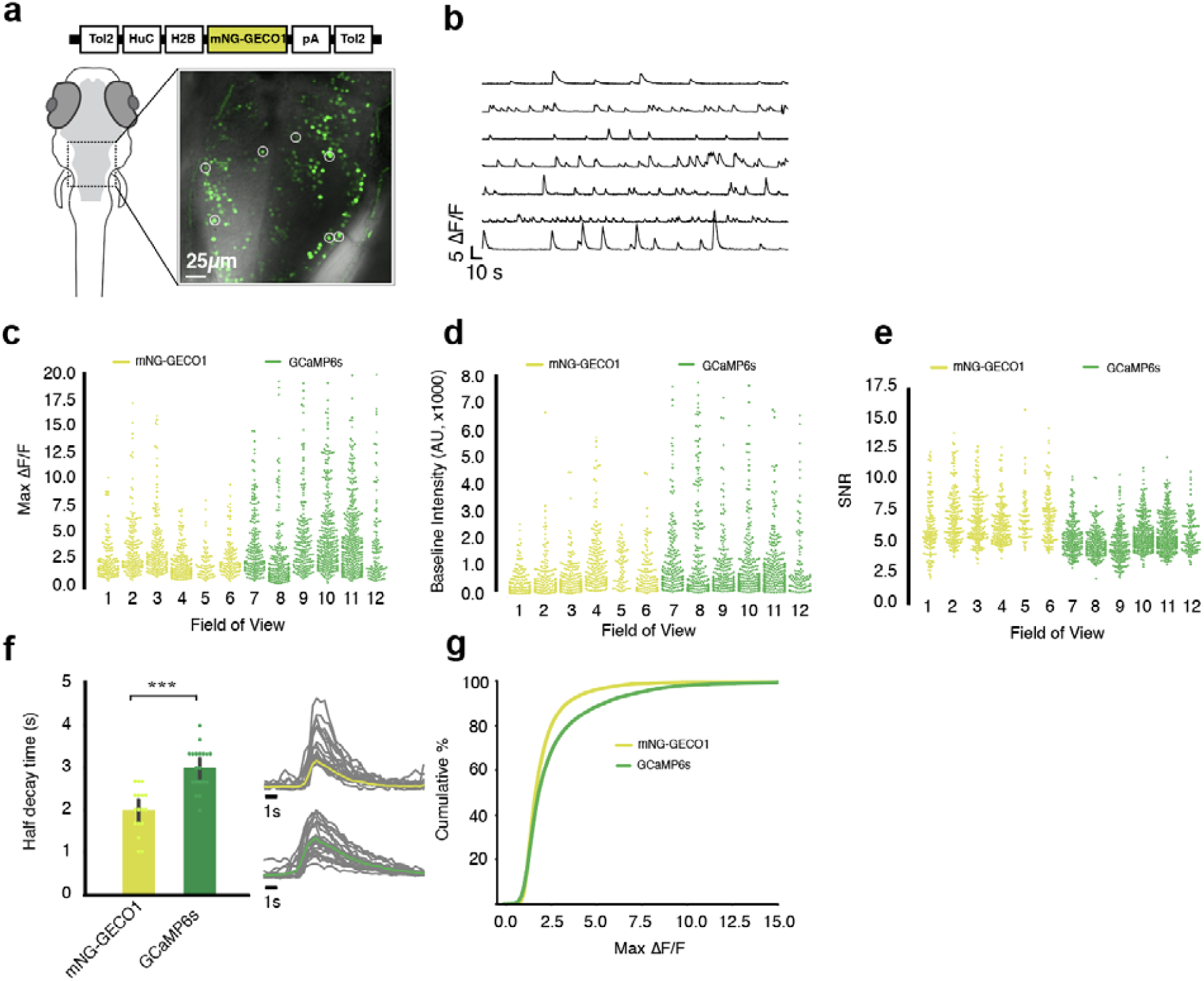
Characterization of mNG-GECO1 and GCaMP6s in transgenic zebrafish hind brain tissue. **a** Schematic representation of Tol2[HuC-H2B-mNG-GECO1] construct and confocal image of one fish (5 to 6 days post fertilization) with 7 region of interests (ROI) circled. **b** Traces of ROI’s from a). **c** Max ΔF/F_0_ calculated by taking the max peak of each cell within the field of interest over 5 minutes; six ROI’s each are used from 5 independent fish expressing mNG-GECO1 and 5 fish expressing GCaMP6s. **d** Baseline fluorescence intensity of each cell within all ROI’s from 5 fish; confocal settings are kept consistent between GCaMP6s and mNG-GECO1 imaging. **e** Signal-to-noise ratio (SNR) computed by dividing ΔF/F_0_ by raw standard deviation of each cell across 6 FOV’s each for both sensors. **f** Average half decay time plotted for mNG-GECO1 (n = 17) and GCaMP6s (n = 19) by averaging randomly selected peaks. **g** Cumulative distribution of mNG-GECO1 vs. GCaMP6s. All cells are arranged in incremental order of ΔF/F_0_ and plotted with respect to their ΔF/F_0_ and their position in the order (%).

### Ca^2+^ imaging in human iPSC-derived cardiomyocytes

Chemical Ca^2+^ dyes such as Fluo-4 acetoxymethyl (AM), Rhod-2 AM and Fura-2 AM are often used to phenotype Ca^2+^ transients in induced pluripotent stem cell-derived cardiomyocytes (iSPC-CM). However, these dyes can be toxic^28, 29^ and may potentially suppress the activity of Na^+^ and K^+^-dependent adenosine triphosphatase^30^. As such, we tested whether mNG-GECO1 could serve as a robust tool for observing cells signaling and drug response while preventing cellular toxicity in iPSC-CMs **(Supplementary Fig. 7).** We found that when iPSC-CMs expressing mNG-GECO1 o Fluo-4 AM were treated with 20 mM caffeine, mNG-GECO1 had a 2.8 fold higher ΔF/F response than Fluo-4 (ΔF/F = 11.77 ± 2.82 and 4.18 ± 1.27, respectively) **(Supplementary Fig. 7a, b).** However, when cells were subjected to 0.33 Hz electrical stimulation for 30 minutes, mNG-GECO1 had a slightly lower peak ΔF/F (2.26 ± 0.81) than Fluo-4 AM (3.31 ± 1.42) **(Supplementary Fig. 7c, d).** We suspect that this discrepancy is due to mNG-GECO1s lower affinity for Ca^2+^. When we stimulated the cells in the presence of 20 mM caffeine, the max ΔF/F of mNG-GECO1 (ΔF/F = 14.20 ± 4.67) was higher than the max ΔF/F of Fluo-4 AM (ΔF/F = 6.73 ± 1.19) **(Supplementary Fig. 7e, f).** Based on this data, we propose that mNG-GECO1 may serve as a useful tool for phenotypic screening and functional tests in iPSC-CMs.

## Conclusion

mNG-GECO1 is a new, first-generation, genetically encodable Ca^2+^ indicator that provides performance comparable to 6^th^ and 7^th^ generation GCaMP indicators. We have demonstrated that the *in vitro* performance of mNG-GECO1 in cultured HeLa cells is on par or better than GCaMP6s. However, *in vitro* cultured neuron benchmarking as well as *in vivo* imaging in transgenic zebrafish larvae have indicated that further directed evolution efforts will be required to produce an mNG-GECO1 variant that provides substantial advantages relative to the jGCaMP7 series. Further development of this indicator may come from increasing the Ca^2+^ affinity which would enable more accurate single spike detection. In summary, we have developed a first generation GECI from the mNG scaffold that retains the high fluorescent brightness *in vitro* with performance comparable to the state-of-the-art GECI, GCaMP6s. We expect mNG-GECO1 to be just as amenable to further optimization as the first generation GCaMP, and so mNG-GECO1 is likely to serve as the parent of a new and improved lineage of high performance GECIs.

## Author contributions

LZ, AA, and TP performed the directed evolution experiments and *in vitro* characterization. RP performed the *in vitro* 2P characterization, IK and TGP conducted and analyzed the cultured neuron experiments. RK conducted the initial expression experiments of mNG-GECO1 and its variants in zebrafish under the supervision of WTA. VR completed the zebrafish characterization under the supervision of MA. HYH and YFC did the experiments in human iPSC-derived cardiomyocytes. LZ, AA, KP, and REC wrote and edited the manuscript. All authors were allowed to review and edit the manuscript before publication.

## Acknowledgments

We thank the University of Alberta Molecular Biology Services Unit (Alberta) and Molecular Bio (Janelia) for technical support. We thank Christopher Cairo (Alberta), Andy Holt (Alberta) and Loren Looger (Janelia) for providing access to the instrumentation, Eric Schreiter (Janelia) for providing access to resources and for useful feedback regarding the manuscript. We thank Deepika Walpia (Janelia) and the JRC Histology group for preparing cultured neurons. We thank John Macklin (Janelia) for overseeing two-photon measurements. We thank Nathan Shaner for the mNeonGreen fluorescent protein. REC acknowledges the Japan Society for the Promotion of Science (JSPS), Natural Sciences and Engineering Research Council of Canada (NSERC), and Canadian Institutes of Health Research (CIHR), for funding support. The mNG gene was a kind gift from Jiwu Wang at Allele Biotech.

## Methods

### General procedures

Synthetic DNA oligonucleotides and gBlocks were purchased from Integrated DNA Technologies. Plastic consumables, restriction endonucleases, Taq polymerase, Phusion polymerase, T4 DNA ligase, deoxynucleotides, DH10B *E. coli*, pBAD/His B plasmid, pcDNA3.1(+) plasmid, Bacterial Protein Extraction Reagent (B-PER), Penicillin-Streptomycin, Fetal Bovine Serum (FBS), TurboFect, Lipofectamine 2000, and GeneJet gel or plasmid purification kits were purchased from Thermo Scientific. Endotoxin-free plasmid DNA isolation kits were purchased from Qiagen (cat. 12362). Agarose, MnCl_2_ · 4H_2_O tryptone, D-glucose, ampicillin, L-arabinose, Hank’s balanced salt solution (HBSS), DMEM, TrypLE Express, and LB Lennox media were purchased from Fisher Scientific. NbActiv4 and neuron transfection media were purchased from Brain Bits.

3-(N-morpholino)propanesulfonic acid (MOPS), ethylene glycol-bis(2-aminoethylether)-N,N,N′,N′-tetraacetic acid (EGTA), and nitrilotriacetic acid (NTA), were purchased from VWR. Nickel NTA immobilized metal affinity chromatography protein purification beads were purchased from G-BioSciences. Ionomycin and tricaine methanesulfonate were purchased from Millipore-Sigma. Ethidium bromide and PCR machines (T100 Thermal Cycler) were purchased from BioRad. Gibson Assembly reagent was purchased from New England Biolabs (NEB). Genemorph II Random Mutagenesis kits and QuikChange mutagenesis kits were purchased from Agilent Technologies. Nunc 96-Well Polypropylene DeepWell Storage Plates (cat. 278743) and 96-well Nunc MicroWell 96-Well Optical-Bottom Plates (cat. 265301) were purchased from Thermo Scientific. Molecular weight cut off filters were purchased from Millipore-Sigma. Sequencing was completed by the Molecular Biology Services Unit at the University of Alberta.

### Molecular biology and protein engineering

Libraries for iterative directed evolution were created using Genemorph II Random Mutagenesis kits and NEB’s Gibson Assembly reagent. Blunt ended linear DNA fragments with random mutations are created using the Genemorph II kit according to the manufacturer’s recommendations. Genemorph II PCR product was ligated using NEB Gibson Assembly reagent into a linearized pBAD vector cut with XhoI/HindIII. Site saturation mutagenesis libraries were created using single and multi QuikChange mutagenesis kits according the manufacturers recommendations.

Libraries are transformed into DH10B *E. coli* and plated on 100 μg/L ampicillin/1.5% agar plates with 0.02% L-arabinose and grown overnight (12-18 hours) at 37 °C. Colonies are selected on the basis of fluorescence intensity, picked, and placed into 96 DeepWell blocks containing 1.3 mL of LB Lennox media supplemented with 100 μg/mL ampicillin and 0.02% L-arabinose. Deepwell blocks were shaken overnight at 37 °C. The next day, blocks are centrifuged at 6000 × g for 5 minutes to pellet cells. Media was discarded and 30 μL of B-PER was added to each well. After shaking for 15 minutes, 200 μL of 10 mM EGTA in 30 mM MOPS/100 mM KCl pH 7.2 (MOPS/KCl buffer) is added to each well of the blocks before mixing briefly and being centrifuged again for 5 minutes at 6000 × g. 90 μL of the resulting lysate is loaded in each well of 96-well optical bottom plates. Fluorescence intensity for each well of the plate is then read with a Tecan Safire^2^ microplate reader to determine the low Ca^2+^ intensity for each variant. High Ca^2+^ intensity is acquired by adding 15 μL of 100 mM Ca^2+^ in 30 mM MOPS pH 7.2 with a 60 second shake before reading. Taking the value of the high Ca^2+^ intensity divided by the low Ca^2+^ intensity gives a relative sensitivity value. Promising candidates, usually 10% of each 96-well block, are retested from the lysate in 10 mM low (EGTA chelated) and 10 mM high Ca^2+^ solution diluted in MOPS/KCl buffer. The plasmids associated with the promising variants are sent for sequencing and used as template for the next round of directed evolution. For cultured neuron field stimulation experiments, GCaMP6s, jGCaMP7f, jGCaMP7s, jGCaMP7c, and jGCaMP7b plasmids (available on Addgene) were subcloned into a syn-<*GCaMP*>-IRES-mCherry-WPRE-pA vector.

Constructs for zebrafish transfection were created by ligating mNG-GECO1 PCR product into a Tol2 transposon backbone. Briefly, PCR of mNG-GECO variants were ligated into Tol2-HuC-H2B vector (Addgene plasmid #59530) cut with SalI/AgeI using Gibson Assembly. The ligated constructs were transformed into NEB Turbo Competent *E. coli* cells and grown in 250 μL culture overnight at 30 °C. The next day, the culture is pelleted, and the DNA is purified using endotoxin-free plasmid DNA purification protocol using EndoFree Plasmid Maxi Kit. The DNA is eluted with EF-free H_2_O and verified by sequencing.

### Protein purification and *in vitro* characterization

To purify mNG, mNG-GECO variants, and GCaMP6s for *in vitro* characterization, pBAD/His B plasmid containing the gene of interest was used to transform electrocompetent DH10B *E. coli*, which were then streaked on 100 μg/mL ampicillin/1.5% agar plates. After overnight incubation at 37 °C, a single colony was picked and inoculated to a 2 L flask containing 500 mL of 100 μg/mL ampicillin/0.02% L-arabinose liquid media and cultured for 24-30 hours at 37 °C. The culture is then centrifuged at 6000 × g for 6 minutes to collect the cells. Cells are re-suspended in 30 mL of cold Tris buffered saline (TBS, 150 mM NaCl, 50 mM Tris-HCl) pH 8.0 and lysed by sonication (QSonica Q700, amplitude 50, 1 second on, 2 seconds off, 3 minutes sonication time). All subsequent purification procedures were performed on ice. The resulting lysate was clarified of cell debris by centrifugation for 1 hour at 21,000 × g, filtered through a Kim-wipe into a 50 mL conical bottom tube, and incubated for 3 hours with Ni-NTA resin. Resin containing NTA bound protein was washed with 100 mL of 20 mM imidazole TBS wash buffer and eluted with 250 mM imidazole TBS elution buffer. Purified protein was buffer exchanged into TBS using a 10,000 Da molecular weight cut-off filter (Millipore-Sigma) through 3 successive washes. Absorption spectra were recorded on a Beckman-Coulter DU-800 UV-visible spectrophotometer and fluorescence spectra recorded on a Tecan Safire^2^ plate reader.

Extinction coefficient determination for mNG-GECO variants were performed using the alkaline denaturation method with mNG as a standard^31^. Briefly, the concentration of protein was adjusted by dilution in MOPS/KCl pH 7.2 to reach an absorbance of 0.6 to 1.0. A dilution series with MOPS/KCl and 10 mM Ca^2+^ was then prepared with absorbances of 0.01, 0.02, 0.03, 0.04, and 0.05 for mNG, mNG-GECO variants, and GCaMP6s. Integration of the fluorescent peaks provides a total fluorescent emission value which was plotted against the absorbance to provide a slope. The quantum yields of mNG-GECO variants were determined using the published^31^ QY value of mNG in a ratiometric manner:

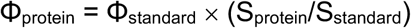

Extinction coefficients were determined by measuring the absorption spectrum in MOPS/KCl pH 7.2 and 2 M NaOH. The absorbance value for the denatured GFP peak at 440 nm was divided by the previously determined extinction coefficient of 44,000 M^−1^cm^−1^ to give the concentration of protein^31^. Using Beer’s law, the extinction coefficient was then determined by dividing the TBS sample absorbance maximum by the calculated protein concentration.

Determination of *K*_d_ was performed as previously described^32, 33^. Briefly, a reciprocal dilution series was created with either 10 mM EGTA/10 mM Ca^2+^ EGTA ranging in free Ca^2+^ concentration of 0 to 0.039 mM or 10 mM NTA/10 mM Ca^2+^ NTA ranging in free Ca^2+^ concentration from 0 to 1.13 mM^33^. An equal amount of purified mNG-GECO was diluted 100× into 100 μL of buffer and the intensity plotted against free Ca^2+^ in triplicate. The data are then fit to a four-parameter variable-slope in GraphPad Prism 7 software to determine the *K*_d_.

### Two Photon Measurements

The two photon measurements were performed in 39 μM free Ca^2+^ (+Ca^2+^) buffer (30 mM MOPS, 10 mM CaEGTA in 100 mM KCl, pH 7.2) or 0 μM free Ca^2+^ (−Ca^2+^) buffer (30 mM MOPS, 10 mM EGTA in 100 mM KCl, pH 7.2). The two photon excitation spectra were acquired as previously described^2^. Protein solution of 2 – 4 μM concentration in +Ca^2+^ or −Ca^2+^ buffer was prepared and measured using an inverted microscope (IX81, Olympus) equipped with a 60×, 1.2 NA water immersion objective (Olympus). Two photon excitation was obtained using an 80 MHz Ti-Sapphire laser (Chameleon Ultra II, Coherent) for spectra from 710 nm to 1080 nm. Fluorescence collected by the objective was passed through a short pass filter (720SP, Semrock) and a band pass filter (550BP200, Semrock), and detected by a fiber-coupled Avalanche Photodiode (APD) (SPCM_AQRH-14, Perkin Elmer). The obtained two photon excitation spectra were normalized for 1 μM concentration and further used to obtain the action cross-section spectra (AXS) with fluorescein as a reference^34^.

Fluorescence correlation spectroscopy (FCS) was used to obtain the 2P molecular brightness of the protein molecule. The molecular brightness was defined by the rate of fluorescence obtained per total number of emitting molecules. 50-200 nM protein solutions were prepared in +Ca^2+^ buffer and excited with 940 nm wavelength at various power ranging from 2-30 mW for 200 seconds. The obtained fluorescence was collected by an APD and fed to an autocorrelator (Flex03LQ, Correlator.com). The obtained autocorrelation curve was fit on a diffusion model through an inbuilt Matlab function^35^ to determine the number of molecules <N> present in the focal volume. The 2-photon molecular brightness (*ε*) at each laser power was calculated as the average rate of fluorescence <*F*> per emitting molecule <*N*>, defined as *ε* = <*F*>/<*N*> in kilocounts per second per molecule (kcpsm). As a function of laser power, the molecular brightness initially increases with increasing laser power, then levels off and decreases due to photobleaching or saturation of the protein chromophore in the excitation volume. The maximum or peak brightness achieved, <*ε_max_*>, represents a proxy for the photostability of a fluorophore.

### *In vitro* kinetics analysis by stopped-flow

Rapid kinetic measurements of purified mNG-GECO1 and GCaMP6s were made using an Applied Photophysics SX-20 Stopped-flow Reaction Analyzer exciting at 488 nm with 2 nm bandwidth and collecting light at 520 nm through a 10 mm path at room temperature. Briefly, 2 μM of mNG-GECO1 and GCaMP6s proteins in 1 mM Ca^2+^ (30 mM MOPS, 100 mM KCl, pH 7.2) were rapidly mixed at 1:1 ratio with 50 mM of EGTA (same buffer as above) at room temperature. *k*_off_ values were determined by fitting a single exponential dissociation curve to the signal decay using Graphpad Prism, with units of s^−1^. For *k*_on_, both proteins buffered in 30 mM MOPS, 100 mM KCl, 50 μM EGTA were rapidly mixed at 1:1 ratio with varying concentrations of Ca^2+^ produced by reciprocal dilutions of 10 mM EGTA and 10 mM CaEGTA. The measured fluorescence change overtime was fitted using a 2-phase association curve to obtain the slow and fast observed rate constants (*k*_obs_) for each free Ca^2+^ concentration. All measurements were done in triplicates, and values are reported as mean ± s.e.m. where noted.

### Fluorescence live cell imaging

#### Imaging in HeLa cells

We followed previously reported protocols for our Ca^2+^ imaging experiments^36^. Briefly, HeLa cells cultured in DMEM with 10% fetal bovine serum supplemented with penicillin-G potassium salt (50 units/mL) and streptomycin sulphate (50 μg/mL) were plated on collagen coated 35 mm glass bottom dishes. HeLa cells are transfected at 60% confluency with 1 μg of pcDNA3.1(+) harboring the variant of interest using 2 μL of TurboFect according to the manufacturer’s recommendation. After overnight incubation at 37 °C with 5% CO_2_, cells were washed twice with prewarmed Hank’s balanced salt solution immediately before imaging.

Imaging of transfected HeLa cells was performed on an inverted Zeiss 200M microscope with Semrock filters (excitation 470/40, emission 525/50) and captured with an OrcaFlash 4.0 – C13440 (Hamamatsu). Images were acquired through a 40× (N.A. 1.3) oil immersion lens using MetaMorph 7.8.0.0 software and an MS-2000 automated stage (Applied Scientific Instrumentation).

### Imaging in dissociated rat cortical neurons

The mNG-GECO1 indicator was compared to other GECIs in a field stimulation assay^37^. Neonatal (P0) rat pups were euthanized, and their cortices were dissected and dissociated using papain (Worthington). Cells were transfected by combining 5×10^5^ viable cells with 400 ng plasmid DNA and nucleofection solution electroporation cuvettes (Lonza). Electroporation was performed according to the manufacturer instructions. Cells were then plated at a density of 5×10^5^ cells/well in poly-D-lysine (PDL) coated 96-well plates. After 14-18 days in vitro, culture medium was exchanged for an imaging buffer solution with a drug cocktail to inhibit synaptic transmission^37^. The field stimulation assay was performed as previously described^4, 20, 37^. Briefly, neurons were field stimulated (1, 2, 3, 5, 10, 20, 40, 160 pulses at 83 Hz, 1 ms, 40V), and concurrently imaged with an electron multiplying charge coupled device (EMCCD) camera (Andor iXon DU897-BV, 198 Hz, 4×4 binning, 800 × 800 μm, 1,400 frames). Reference images were taken after stimulation to perform cell segmentation during analysis. Illumination was delivered by blue light (470 nm, Cairn Research Ltd; excitation: 450-490 nm; emission: 500-550 nm; dichroic: 495 nm long-pass). The illumination power density was measured to be 19 mW/mm^2^ at the sample. Stimulation pulses were synchronized with the camera using data acquisition cards (National Instruments), controlled with Wavesurfer software (https://wavesurfer.janelia.org/). Imaging was performed at room temperature. Data were analyzed using previously-developed MATLAB (Mathworks) scripts^20, 37^.

### Ca^2+^ imaging in human iPSC-derived cardiomyocytes

Human iPSC-derived cardiomyocytes (human iPSC Cardiomyocytes - male | ax2505) were purchased from Axol Bioscience. The 96 well glass-bottom plate or MatTek glass bottom dish (Ashland, MA, US) were first coated with Fibronectin/Gelatin (0.5% / 0.1%) at 37 °C for at least 1 hour. The cells were plated and cultured for three days in Axol’s Cardiomyocyte Maintenance Medium. The cells then were ready for final observation with Tyrode’s buffer. For electrical stimulations, iPSC derived cardiomyocytes were plated on MatTek glass bottom dish (Ashland, MA, US) at 100,000 cells/well. Electrical stimulation was done with 10 V, 10 ms duration and 3 seconds internal using myopacer (Ion optix c-pace ep). To image, an inverted microscope (DMi8, Leica) equipped with a 63× objective lens (NA 1.4) and a multiwavelength LED light source (pE-4000, CoolLED) was used. iPSC derived cardiomyocytes were plated out as above, and then loaded with 5 μM Fluo-4-AM (Thermo-Fisher, UK) at room temperature for 10 minutes, free dye was washed off by media replacement with pre-heated culture media, followed by imaging with iXon EMCCD (Andor) camera using 488 nm LED illumination. The GFP filter set (DS/FF02-485/20-25, T495lpxr dichroic mirror, and ET525/50 emission filter) was used for Fluo-4 and mNG-GECO1 observation.

### Imaging of zebrafish larvae

To demonstrate the sensitivity and brightness of mNG-GECO1 *in vivo*, we performed fluorescence imaging of Ca^2+^ activity in a subset of neurons in larval zebrafish. Initially, we used the AB/WIK zebrafish strain for morphology studies (**Supplementary Fig. 4)**, which were treated with 1-phenol-2-thiourea (PTU) to inhibit pigmentation, as described previously^38^. Later, *Casper* strains were available and 20 ng/μL DNA plasmids encoding mNG-GECO1 under the control of nuclear-localized elavl3/HuC promoter (Addgene: 59530) were injected into two-cell stage embryos of *Casper* mutant zebrafish^33^ with 40 ng/μl Tol2 transposase mRNA (26) to generate F0 transgenic zebrafish. Imaging experiments were performed using 6 day old embryos. Embryos showing expression were treated with 1 mg/mL bath-applied α-bungarotoxin (Thermo Fischer Scientific, B1601) dissolved in external solution for 30 seconds to block movement, and subsequently incubated with 80 mM 4-aminopyridine (4AP) for 10 minutes. After incubation, the larvae were embedded in 2% low melting temperature agarose to prevent motion. For earlier imaging (**Supplementary Fig. 4**) a Zeiss 700 confocal microscope was used with A-Plan 10×/0.25 Ph1 M27 objective lens to obtain picture from the whole larvae (**Supplementary Fig. 4a**). For enlarged areas (**Supplementary Fig. 4b-e**), a Plan-Apochromat 20×/0.8 M27 lens was used. Later imaging was performed using a 488 nm laser (0.45 μM) and a 525/50 nm emission filter at 3 Hz using Zeiss 880 confocal microscope. The laser power was set to 2.3%, gain 720, and pinhole to 29.3% open. Image acquisition, data registration, segmentation and cell traces were handled using theSuite2p package in Python. All animal procedures were approved by the Institutional Animal Care and Use Committee at the HHMI Janelia Research Campus or by the Animal Care and Use Committee: Biosciences at the University of Alberta.

## Supplementary Figures and Tables

**Supplementary Table 1.**
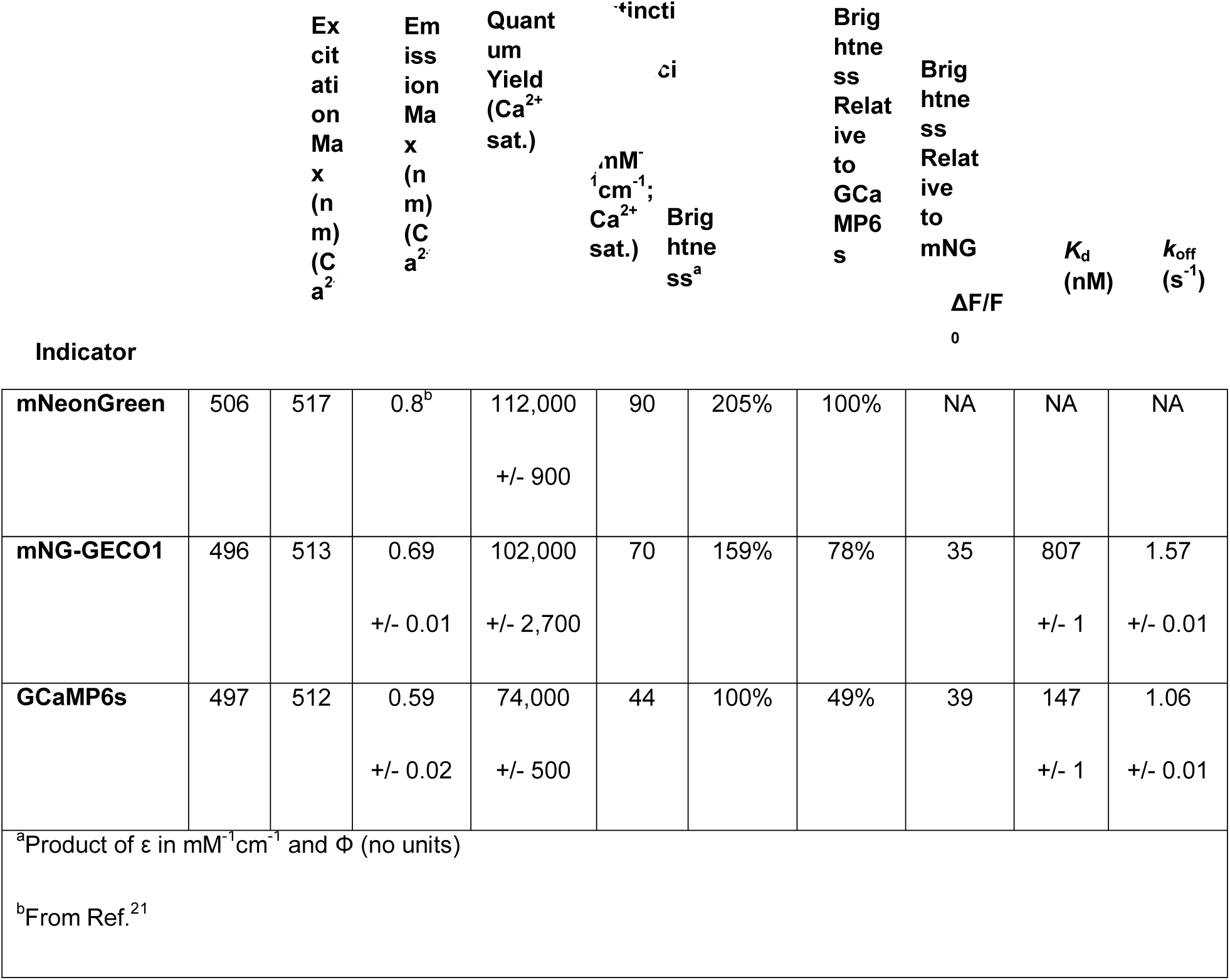
In vitro characterization of mNG-GECO1 and GCaMP6s. mNG, mNG-GECO1, and GCaMP6s were purified and tested in parallel. mNG was used as a standard for brightness and quantum yield determination.

**Supplementary Table 2.** Characterization of Ca^2+^-dependent fluorescence of mNG-GECO1 and GCaMP6s in HeLa cells. Cells were treated with histamine (abb. His), then with Ca^2+^/ionomycin (abb. Ca^2+^), and then with EGTA/ionomycin (abb. EGTA). n is the total number of cells recorded over five independent transfections. The oscillations were detected in all cells with a prominence of greater than 0.5 using a Matlab script. Errors are s.d.

| Protein | Number of HeLa cells (n) | Total Number of oscillations detected | Maximum $\text{Ca}^{2+}$ to minimum EGTA $\Delta F/F_0$ | Maximum His to minimum His ratio | Maximum His to maximum $\text{Ca}^{2+}$ ratio |
| --- | --- | --- | --- | --- | --- |
| mNG-GECO1 | 137 | 1624 | $4.50 \pm 2.96$ | $48.8 \pm 15.1$ | $16.8 \pm 10.5$ |
| GCaMP6s | 99 | 687 | $3.48 \pm 2.40$ | $16.7 \pm 5.2$ | $12.8 \pm 6.11$ |

**Supplementary Table 3.** mNG-GECO1 comparison with GCaMP series sensors in dissociated rat hippocampal neurons. Median values were calculated per well and mean (of medians) ± s.e.m. are presented. SNR values were calculated per individual cells, and median ± s.e.m are shown.

| Protein | 1 AP<br>$\Delta F/F_0$<br>amplitude | 3 AP<br>$\Delta F/F_0$<br>amplitude | 10 AP<br>$\Delta F/F_0$<br>amplitude | 160 AP<br>$\Delta F/F_0$<br>amplitude | Baseline<br>Brightness<br>(Fluorescence<br>Intensity<br>(AU)) | Half<br>rise<br>time 3<br>APs<br>(ms) | Half<br>decay<br>time 3<br>APs (ms) | SNR 1 APs | SNR 3 APs |
| --- | --- | --- | --- | --- | --- | --- | --- | --- | --- |
| mNG-GECO1 | 0.19±0.04 | 0.5±0.07 | 1.5±0.19 | 6.5±0.8 | 1374±31 | 49±1 | 582±12 | 8.9±0.01 | 20.3±0.02 |
| GCaMP6s | 0.27±0.09 | 0.7±0.08 | 3.1±0.26 | 9.0±0.47 | 1302±26 | 65±2 | 1,000±36 | 10.1±0.07 | 25.0±0.03 |
| jGCaMP7b | 0.6±0.07 | 1.2±0.08 | 2.9±0.1 | 3.5±0.09 | 3673±32 | 47±0.1 | 523±2 | 27.4±0.01 | 57.8±0.01 |
| jGCaMP7c | 0.3±0.03 | 0.8±0.04 | 3.9±0.2 | 16.7±0.58 | 770±6 | 49±0.1 | 513±1 | 9.3±0.003 | 21.9±0.01 |
| jGCaMP7f | 0.3±0.06 | 0.8±0.05 | 2.9±0.1 | 7.9±0.25 | 1797±16 | 46±0.1 | 669±2 | 13.6±0.01 | 28.7±0.01 |
| jGCaMP7s | 0.7±0.10 | 2.0±0.11 | 4.6±0.17 | 4.2±0.12 | 1397±11 | 42±0.1 | 736±2 | 24.5±0.01 | 57.9±0.02 |

**Supplementary Table 4.** mNG-GECO1 comparison with GCaMP6s in larval zebrafish 6 dpf. Mean values were calculated for the total number of cells (ROI’s) across six FOV’s in five independent fish. Mean ± s.e.m. are presented. For max ΔF/F_0_, the maximum ΔF/F_0_ from each cell is used. For baseline brightness, the raw intensity of the indicator under the same set of imaging conditions is used. For half decay time, 17 and 19 represented cells were used for mNG-GECO1 and GCaMP6s, respectively. The difference in max ΔF/F_0_ for the two indicators is significant (Kolmogorov-Smirnov statistic = 0.218, p-value = 1.79 × 10^−21^). The difference in baseline brightness for the two indicators is significant (Kolmogorov-Smirnov statistic = 0.100, p-value ≈ 7.31 × 10^−5^). The difference in half decay time for the two indicators is significant (Kolmogorov-Smirnov statistic = 0.666, p-value ≈ 3.00 × 10^−4^).

| Protein | # of biological replicates | # of total field of views (FOV) | # of total cells (N) | Max $\Delta F/F_0$ | Baseline Brightness ($\times 1000$) (AU) | Signal to noise ratio (SNR) | Half decay time ($s^{-1}$) (n) |
| --- | --- | --- | --- | --- | --- | --- | --- |
| mNG-GECO1 | 5 | 6 | 834 | 3.09<br>$\pm 0.08$ | 0.95<br>$\pm 0.03$ | 6.63<br>$\pm 0.07$ | 1.98 ±<br>0.12<br>(17) |
| GCaMP6s | 5 | 6 | 1280 | 4.56<br>$\pm 0.11$ | 1.41<br>$\pm 0.05$ | 5.25<br>$\pm 0.04$ | 3.00 ±<br>0.12<br>(19) |

**Supplementary Figure 1.**
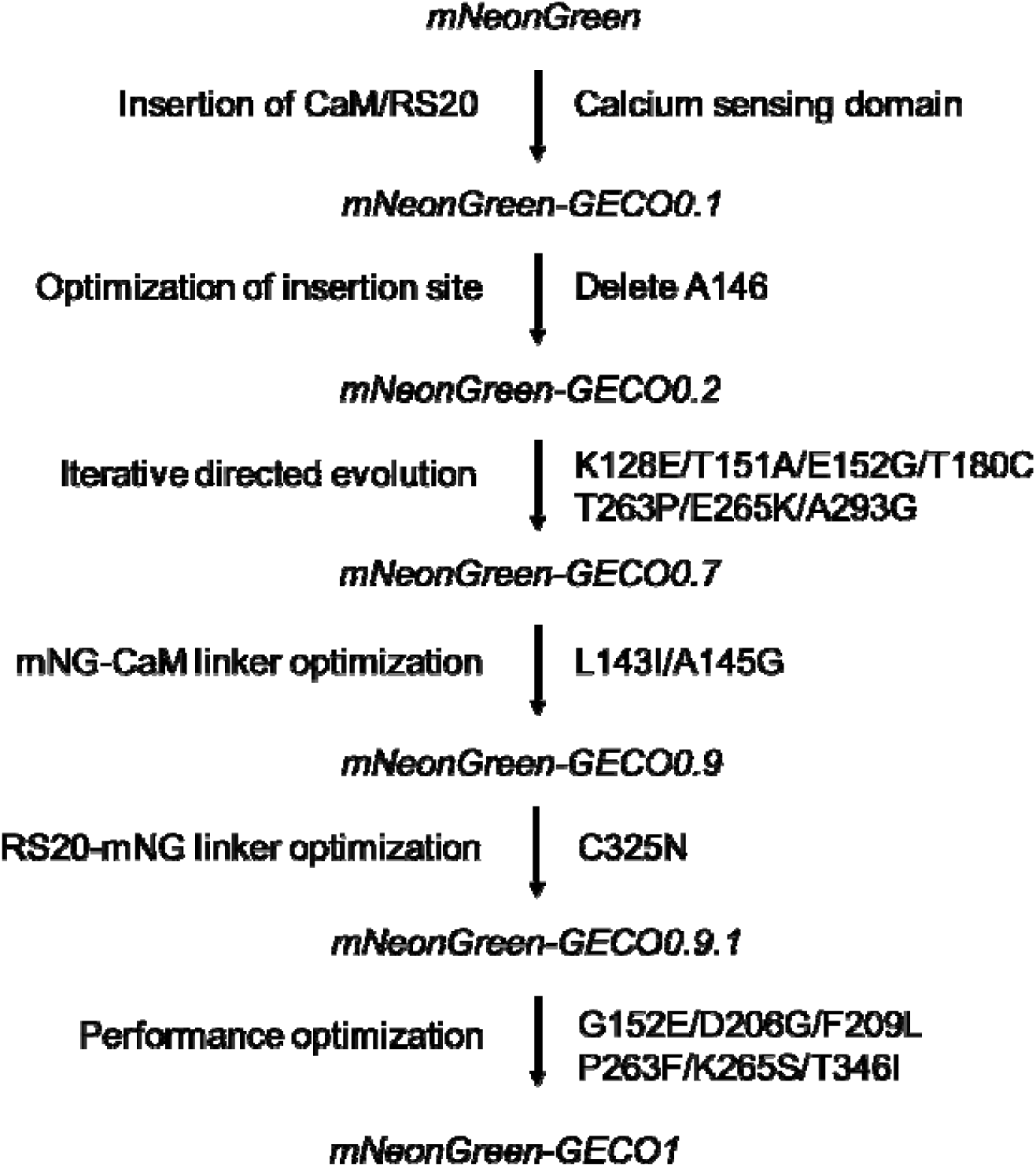
Overview of mNG-GECO1 development. Lineage of mNG-GECO variants starting from initial insertion of Ca^2+^ sensing domain into mNG, and ending with the final mNG-GECO1 variant.

**Supplementary Figure 2.**
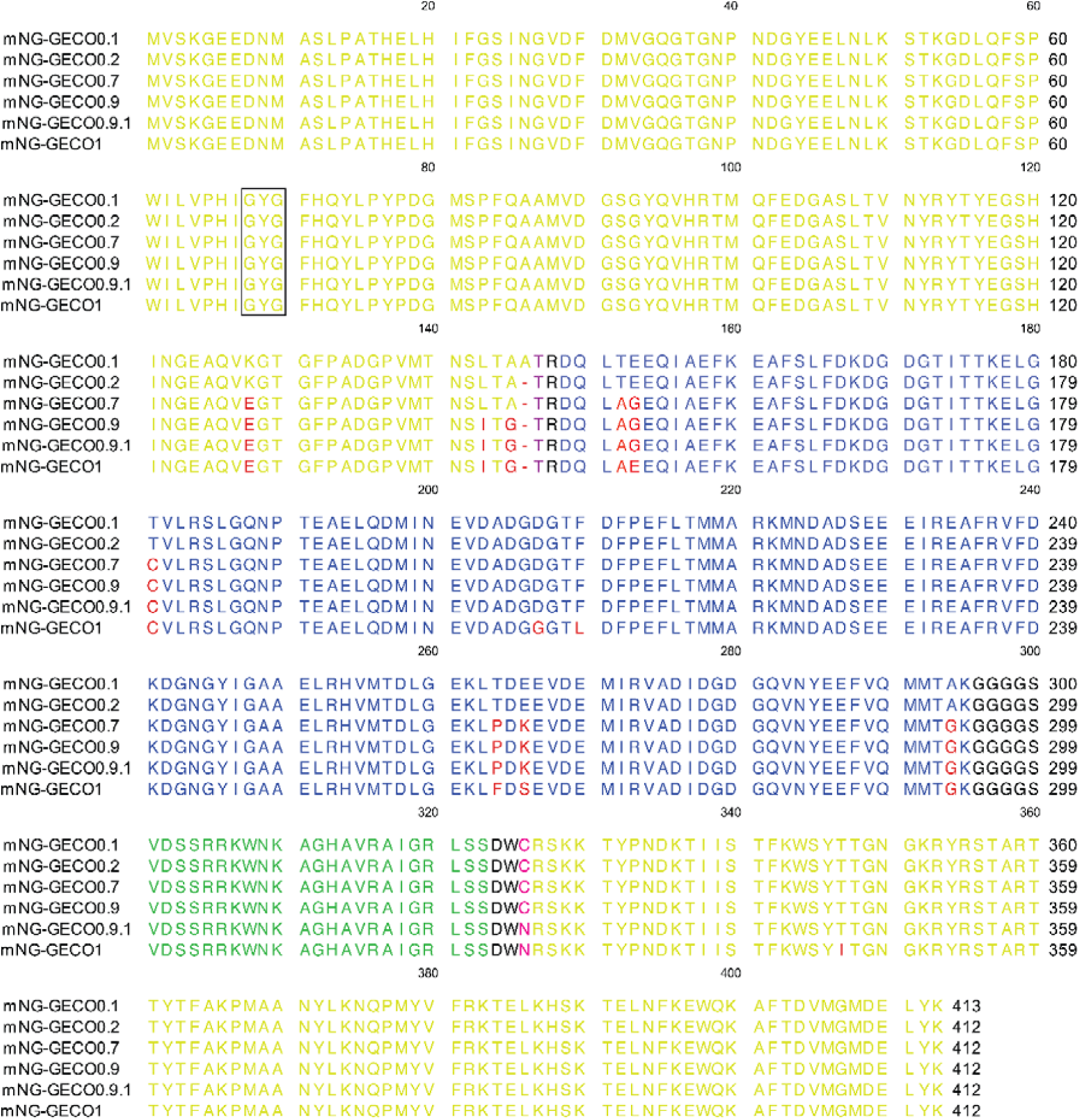
Sequence alignment of mNG-GECO variants. Alignment of mNG-GECO variants 0.2, 0.7, 0.9, 0.9.1, and 1. Similar to the topological representation in **Fig. 1a**, mNG barrel (yellow), CaM (light blue), RS20 (green), linker regions (black), mutations (red), and the chromophore forming residues (black box over “GYG”). Also highlighted are the two residues that flank the insertion site (residue 136 of mNG in purple and residue 139 in magenta; numbering as in PDB ID 5LTR), which are shown as circles in both the protein structure and gene schematic in **Fig. 1a**.

**Supplementary Figure 3.**
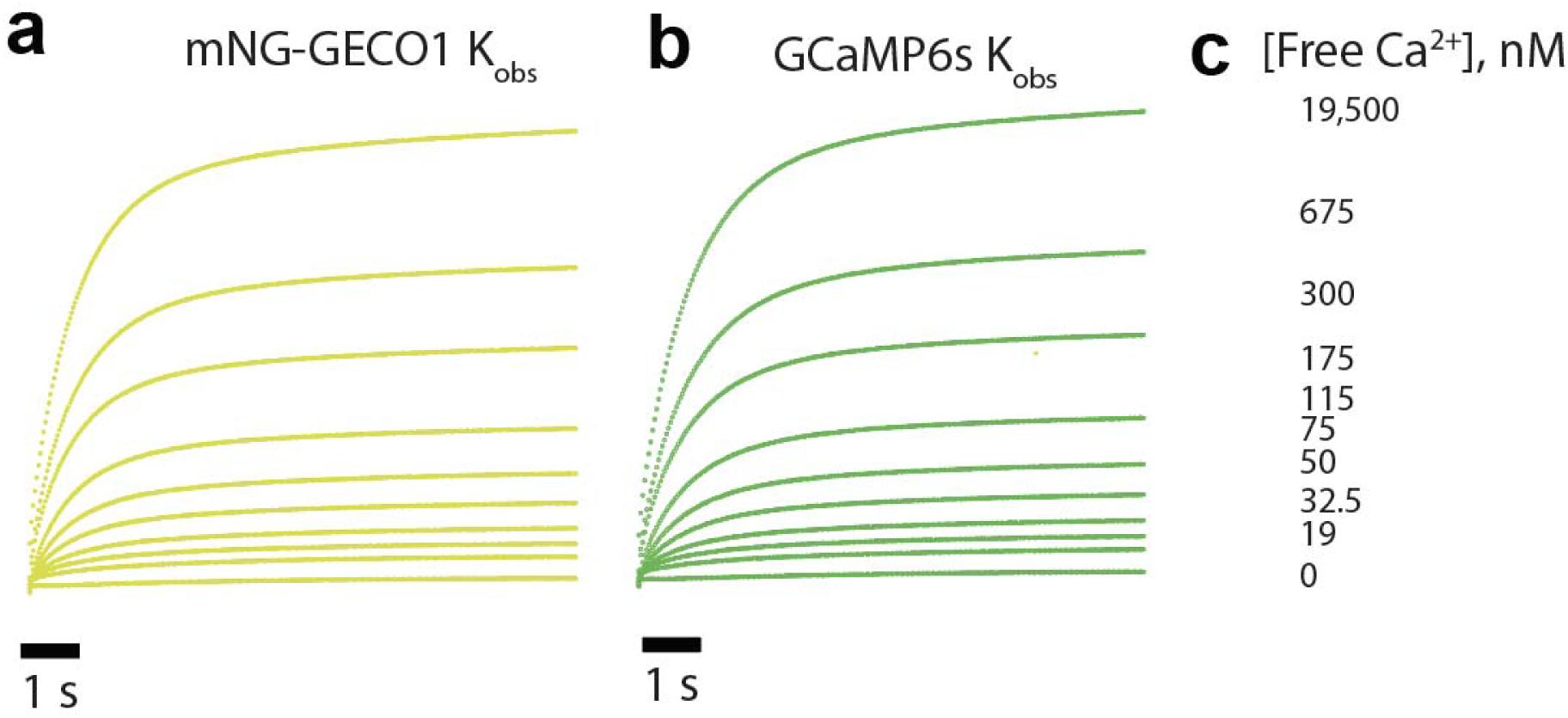
*K*_on_, (observed) traces of mNG-GECO1 and GCaMP6s purified protein using Photophysics SX-20 Stopped-flow. Each protein buffered in 30 mM MOPS, 100 mM KCl, 50 μM EGTA is rapidly mixed at 1:1 ratio with varying concentrations of Ca^2+^ produced by reciprocal dilutions of 10 mM EGTA and 10 mM CaEGTA. **a** mNG-GECO1 change in fluorescence over time as Ca^2+^ is rapidly mixed. **b** GCaMP6s change in fluorescence over time as Ca^2+^ is rapidly mixed. **c** Final free-Ca^2+^ concentrations produced after reciprocal dilutions. mNG-GECO1 and GCaMP6s fit with a double exponential curve due to a slow rate limiting step likely caused by the conformational change of the proteins upon binding to Ca^2+^. Both sensors have a similar t_1/2_ for physiologically relevant Ca^2+^ concentrations.

**Supplementary Figure 4.**
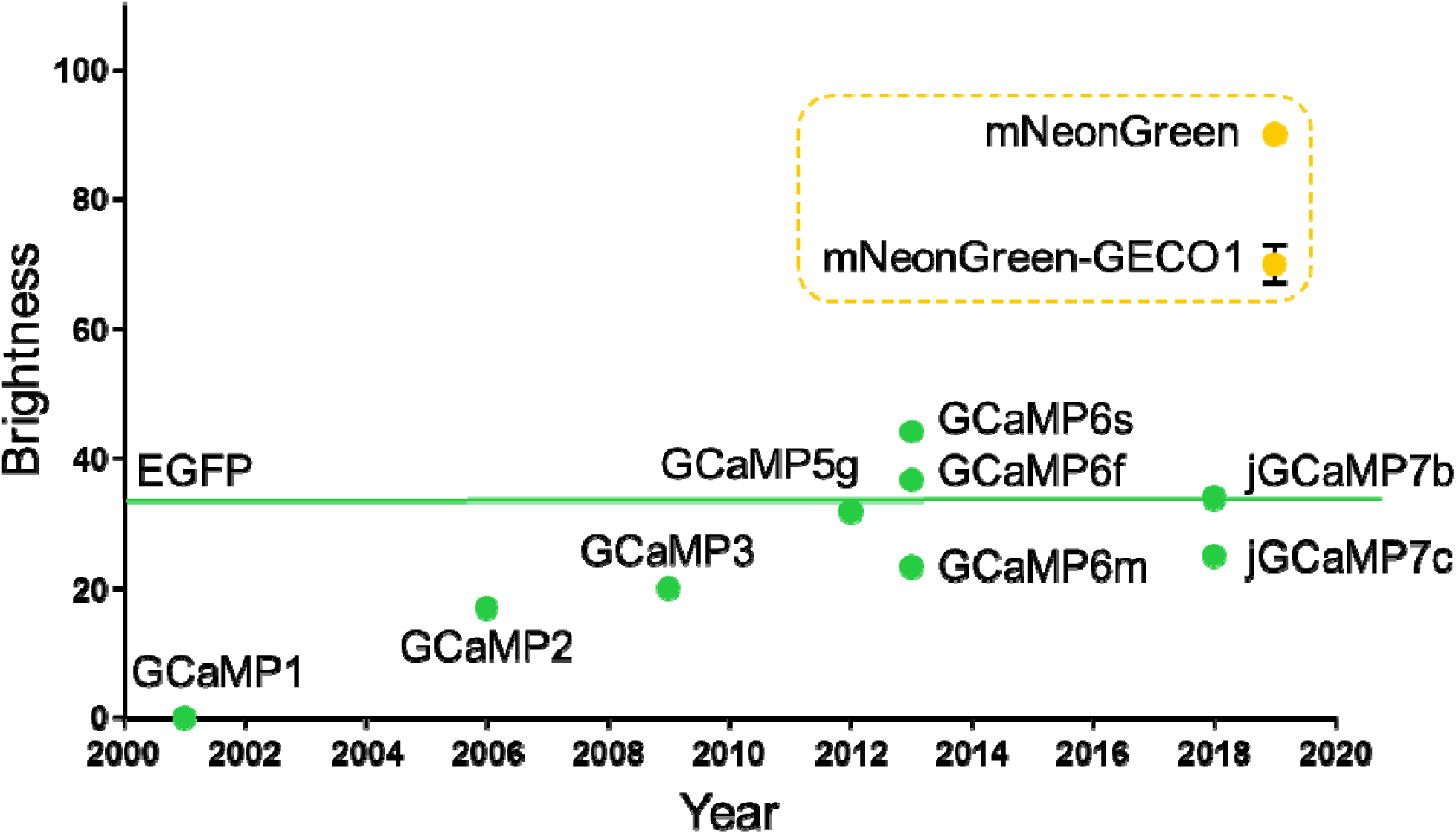
*In vitro* brightness comparison of mNG-GECO1 to GCaMP series. 1P purified protein brightness of first generation mNG-GECO1 compared to the GCaMP series of Ca^2+^ indicators. mNG-GECO1 is substantially brighter *in vitro* than the highly-engineered GCaMP series, which is roughly as bright as its own scaffold, EGFP.

**Supplementary Figure 5.**
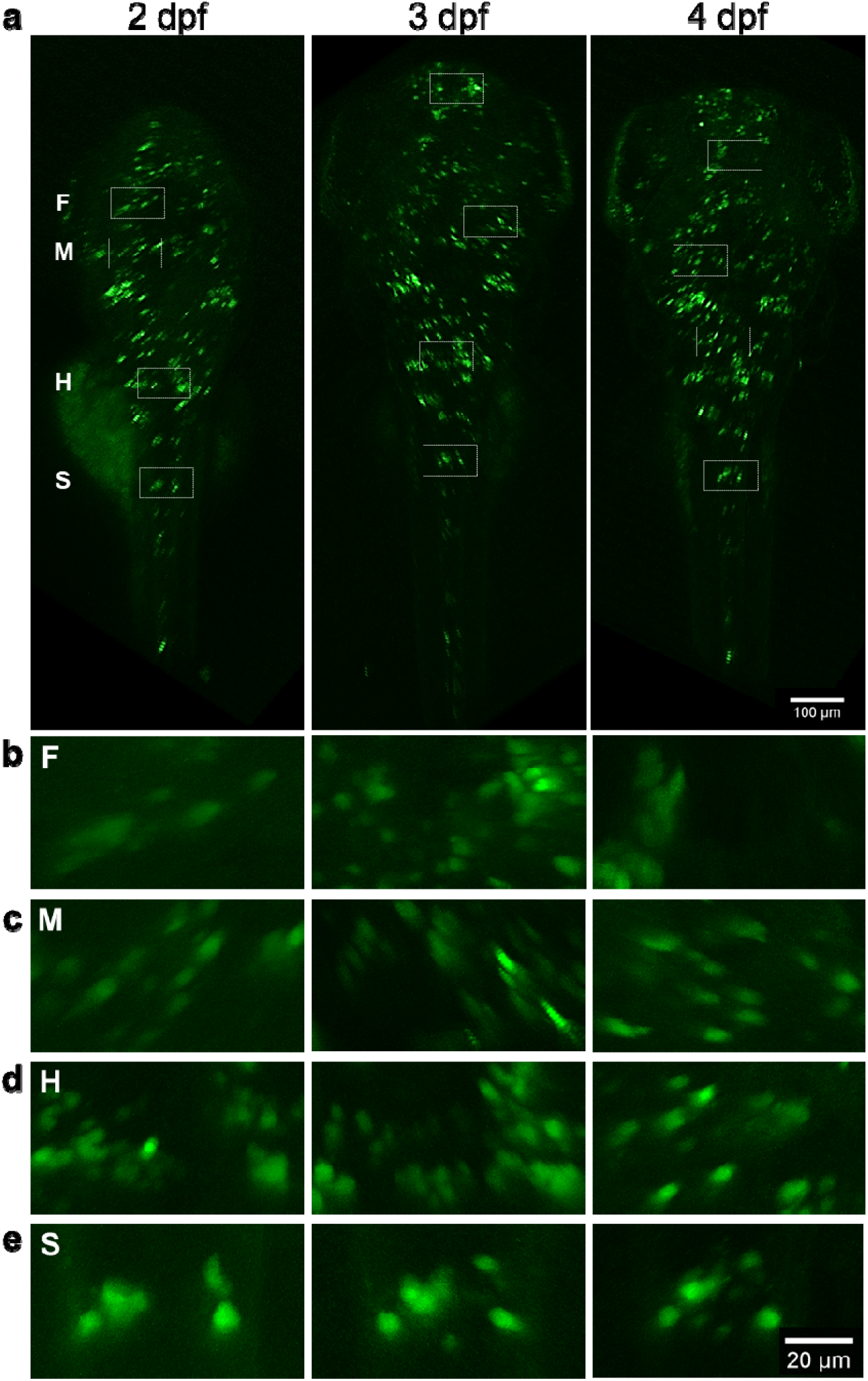
mNG-GECO1 expression profile in zebrafish larvae. Transient expression of mNG-GECO1 in Tg[elavl3:mNG-GECO1] zebrafish at 2, 3 and 4 days post-fertilization. **a** Dorsal view of confocal z-projections obtained from the whole larvae. Small dashed squares mark areas that have been enlarged and presented in b-e. **b-e** Shows areas in the forebrain (F), midbrain (M), hindbrain (H) and spine (S). Scale bars are 100 μm in A and 20 μm in b-e as shown bottom right in e.

**Supplementary Figure 6.**
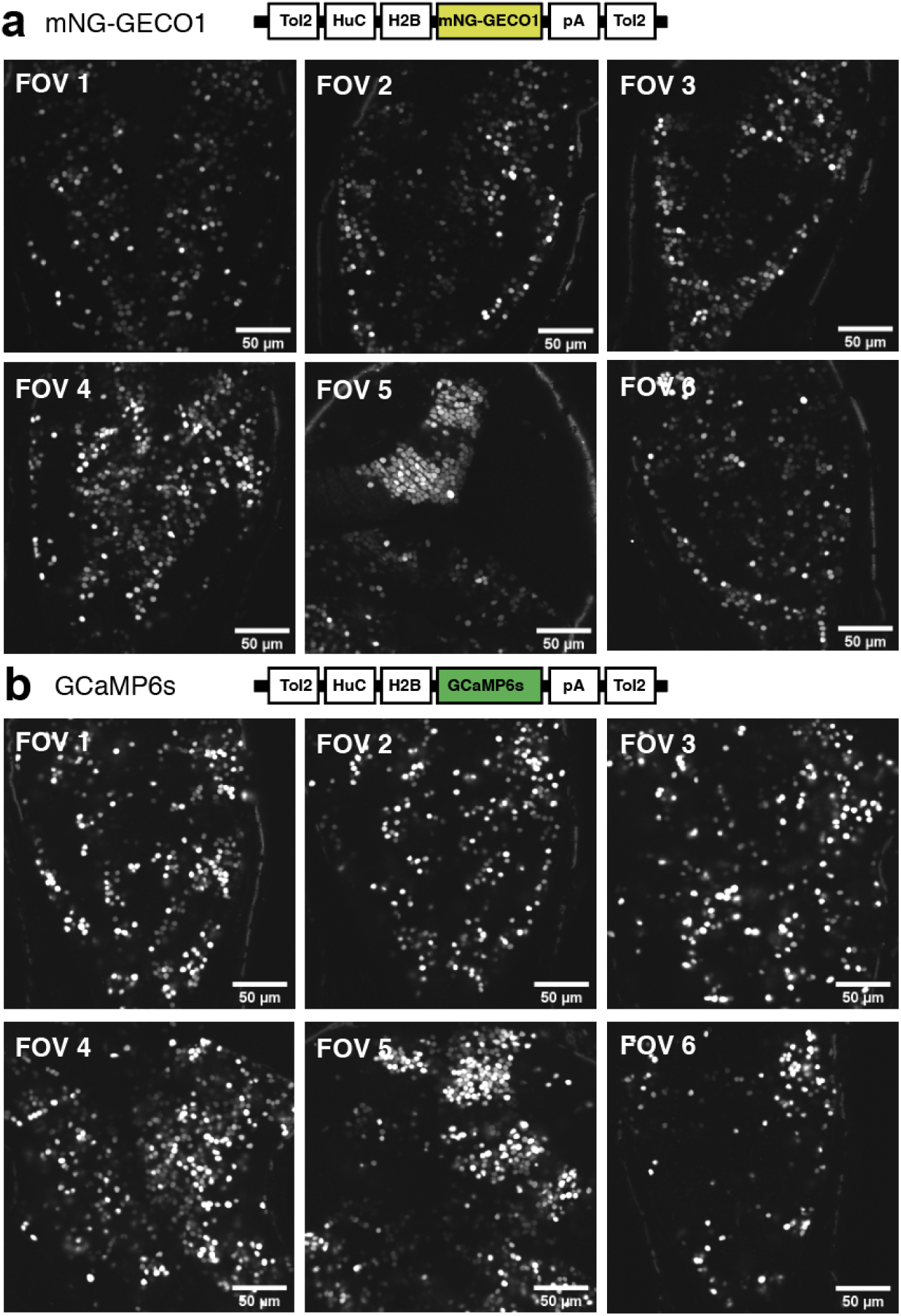
mNG-GECO1 and GCaMP6s field of views (FOV) in zebrafish larvae used for quantification. Transient expression of mNG-GECO1 and GCaMP6s in zebrafish at 6 days post-fertilization. Each FOV image is an average intensity of a 5 minute recording encoding 900 frames. The relative fluorescence intensity is to-scale between the two sensors. **a** FOVs from larvae expressing mNG-GECO1. **b** FOVs from larvae expressing GCaMP6s. Scale bar is 50 μM for all images.

**Supplementary Figure 7.**
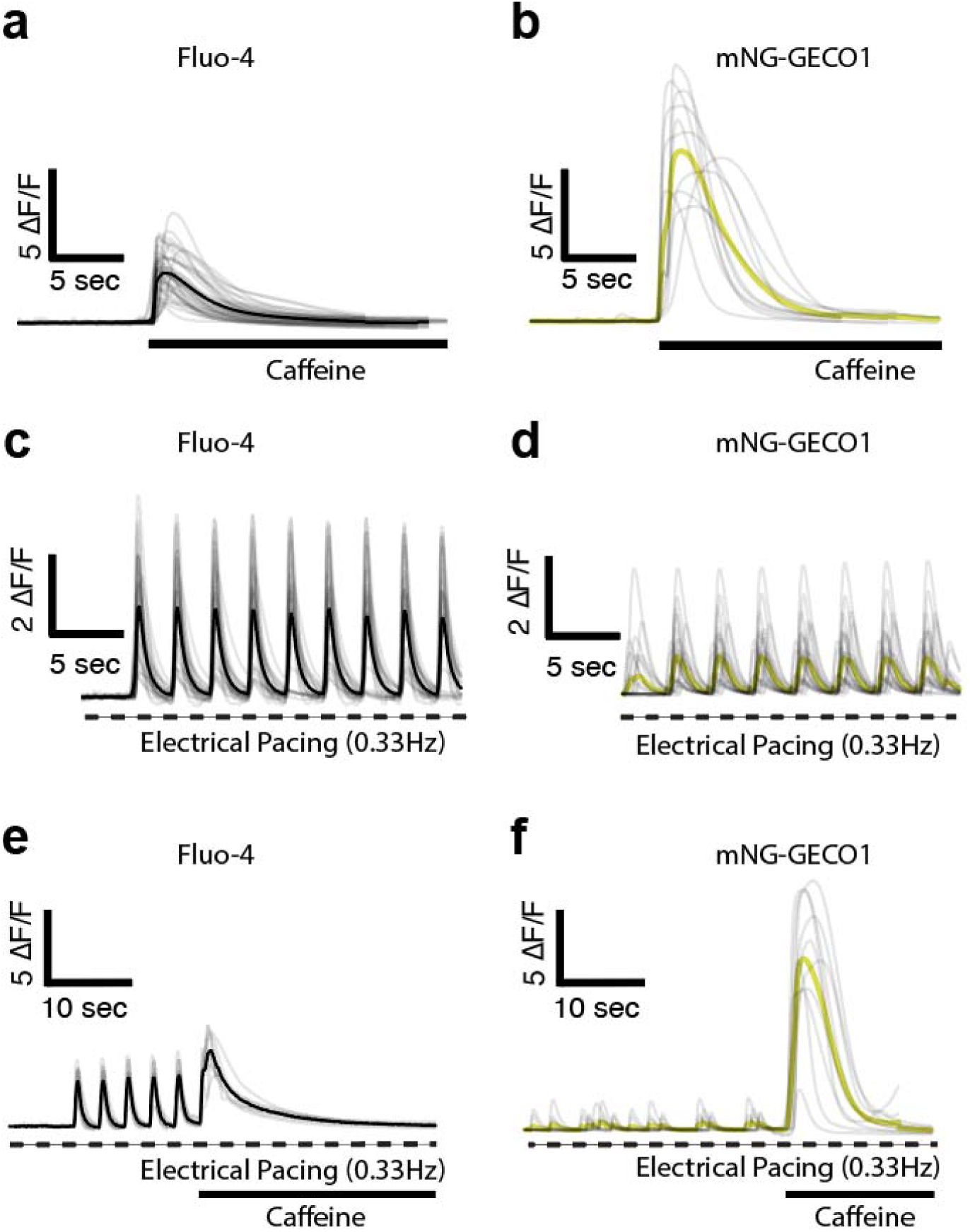
Comparison of Fluo-4 Ca^2+^ dye and mNG-GECO1 in human iPSC-derived cardiomyocytes. **a, b** Single cell Ca^2+^ transient traces from iPSC-CM’s loaded with the Fluo-4 Ca^2+^ dye (n = 38 regions of interests [ROI’s]) or mNG-GECO1 (n=11 ROI’s), respectively. Ca^2+^ transients were evoked using 20 mM caffeine. **c, d** Ca^2+^ transients evoked by electrical pacing were recorded with the Fluo-4 Ca^2+^ dye (n = 25 ROI’s) or mNG-GECO1 (n = 22 ROI’s), respectively. Cells were stimulated electrically at 0.33 Hz and imaged at 10 Hz frame rate. **e, f** Ca^2+^ transients evoked by electrical pacing (0.33 Hz) and caffeine treatment (10 mM) were recorded at 10 Hz with the Fluo-4 Ca^2+^ dye (n = 8 ROI’s) or mNG-GECO1 (n = 8 ROI’s), respectively.

## Notes

#### Summary of Updates

author affiliations updated

